## Supplementary Figures for "In silico screening by AlphaFold2 program revealed the potential binding partners of nuage-localizing proteins and piRNA-related proteins"

### Slide 1
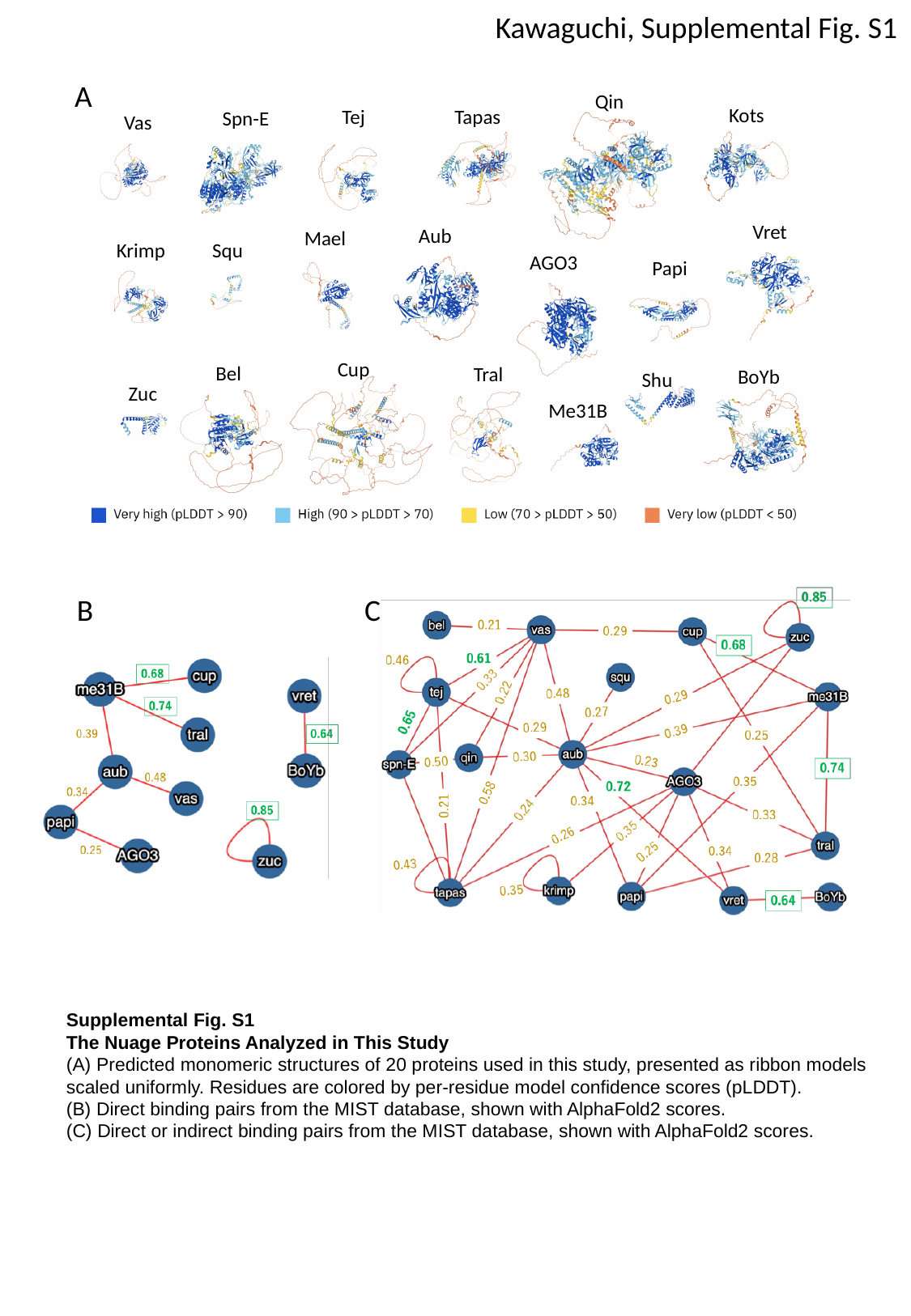

Kawaguchi, Supplemental Fig. S1
A
Qin
Kots
Tej
Tapas
Spn-E
Vas
Vret
Aub
Mael
Krimp
Squ
AGO3
Papi
Cup
Bel
Tral
BoYb
Shu
Zuc
Me31B
B
C
Supplemental Fig. S1
The Nuage Proteins Analyzed in This Study
(A) Predicted monomeric structures of 20 proteins used in this study, presented as ribbon models scaled uniformly. Residues are colored by per-residue model confidence scores (pLDDT).
(B) Direct binding pairs from the MIST database, shown with AlphaFold2 scores.
(C) Direct or indirect binding pairs from the MIST database, shown with AlphaFold2 scores.

### Slide 2
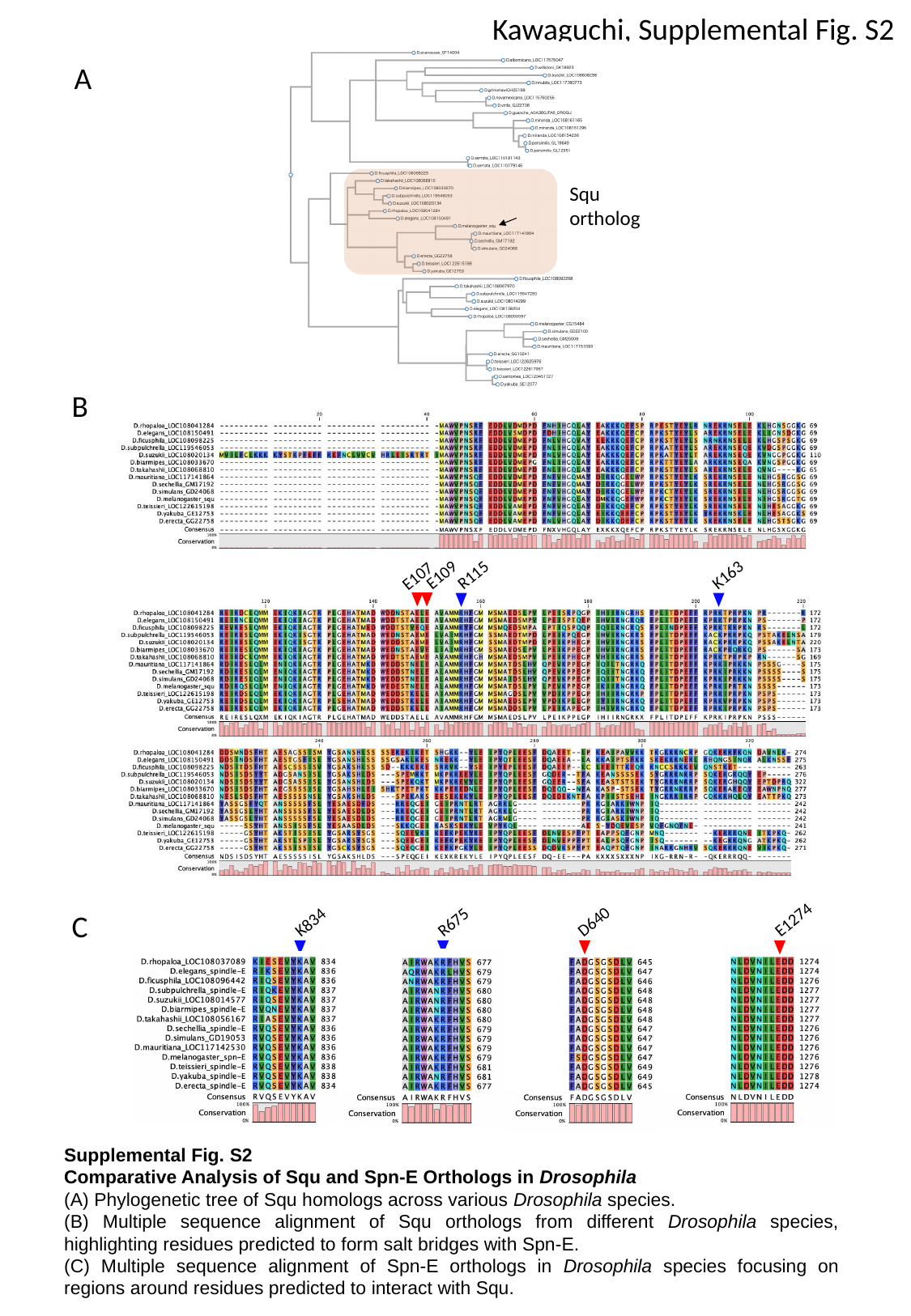

Kawaguchi, Supplemental Fig. S2
Squ
ortholog
A
B
R115
E109
K163
E107
K834
R675
E1274
D640
C
Supplemental Fig. S2
Comparative Analysis of Squ and Spn-E Orthologs in Drosophila
(A) Phylogenetic tree of Squ homologs across various Drosophila species.
(B) Multiple sequence alignment of Squ orthologs from different Drosophila species, highlighting residues predicted to form salt bridges with Spn-E.
(C) Multiple sequence alignment of Spn-E orthologs in Drosophila species focusing on regions around residues predicted to interact with Squ.

### Slide 3
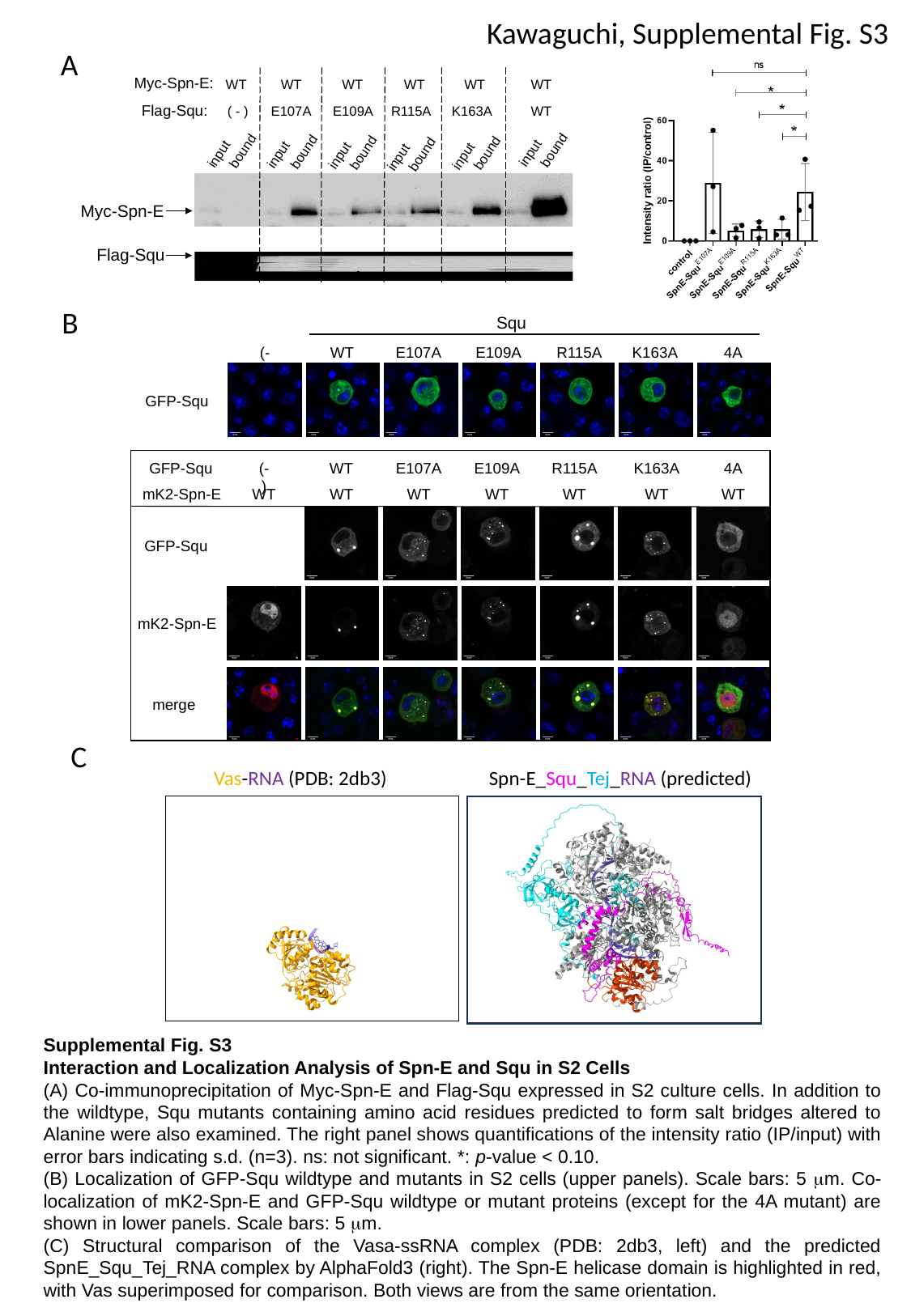

Kawaguchi, Supplemental Fig. S3
A
 Myc-Spn-E:
WT
WT
WT
WT
WT
WT
 Flag-Squ:
( - )
E107A
E109A
R115A
K163A
WT
bound
bound
bound
input
bound
bound
input
input
bound
input
input
input
 Myc-Spn-E
 Flag-Squ
B
Squ
(-)
WT
E107A
E109A
R115A
K163A
4A
GFP-Squ
GFP-Squ
(-)
WT
E107A
E109A
R115A
K163A
4A
mK2-Spn-E
WT
WT
WT
WT
WT
WT
WT
GFP-Squ
mK2-Spn-E
merge
C
Vas-RNA (PDB: 2db3)
Spn-E_Squ_Tej_RNA (predicted)
Supplemental Fig. S3
Interaction and Localization Analysis of Spn-E and Squ in S2 Cells
(A) Co-immunoprecipitation of Myc-Spn-E and Flag-Squ expressed in S2 culture cells. In addition to the wildtype, Squ mutants containing amino acid residues predicted to form salt bridges altered to Alanine were also examined. The right panel shows quantifications of the intensity ratio (IP/input) with error bars indicating s.d. (n=3). ns: not significant. *: p-value < 0.10.
(B) Localization of GFP-Squ wildtype and mutants in S2 cells (upper panels). Scale bars: 5 μm. Co-localization of mK2-Spn-E and GFP-Squ wildtype or mutant proteins (except for the 4A mutant) are shown in lower panels. Scale bars: 5 μm.
(C) Structural comparison of the Vasa-ssRNA complex (PDB: 2db3, left) and the predicted SpnE_Squ_Tej_RNA complex by AlphaFold3 (right). The Spn-E helicase domain is highlighted in red, with Vas superimposed for comparison. Both views are from the same orientation.

### Slide 4
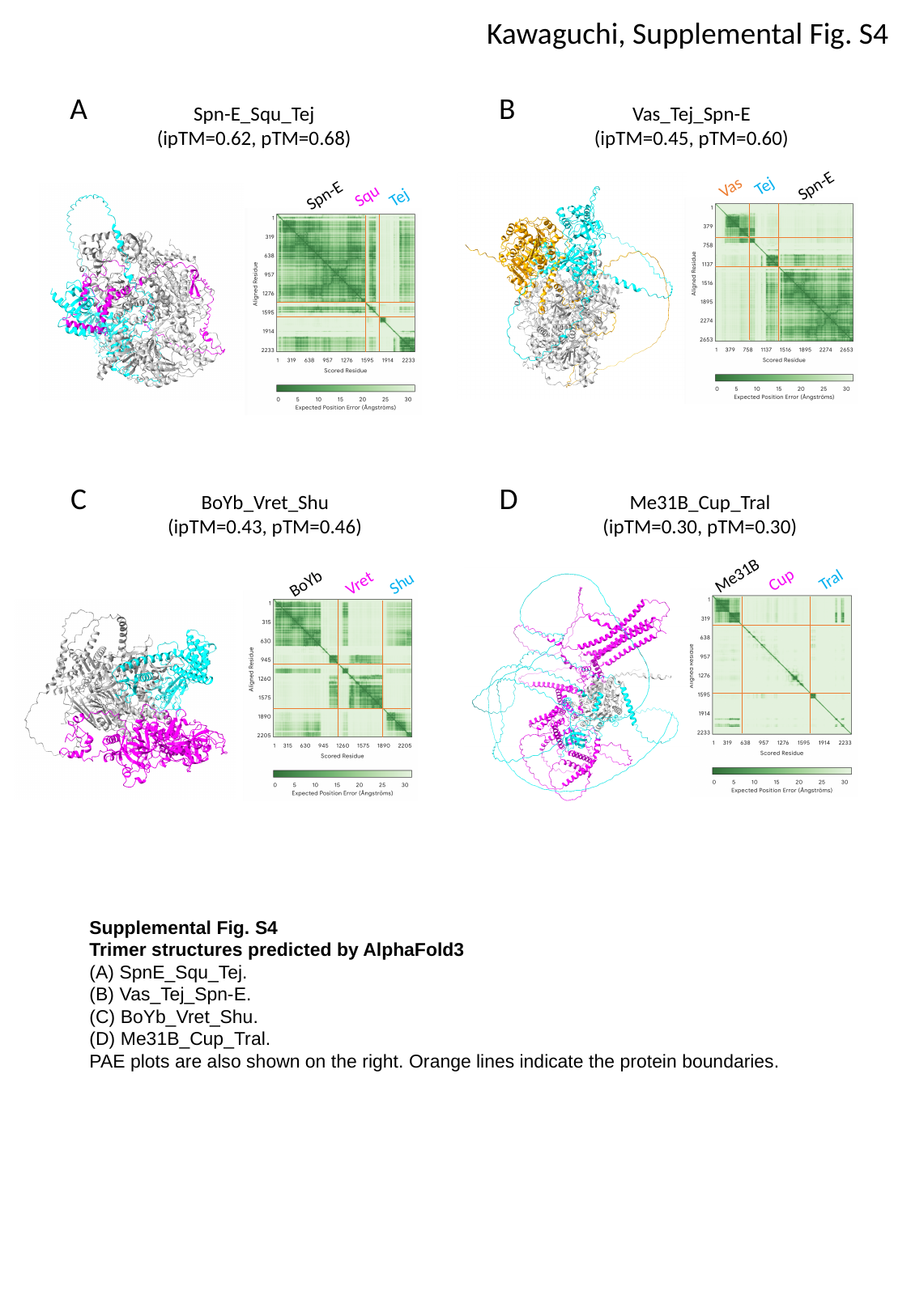

Kawaguchi, Supplemental Fig. S4
A
B
Spn-E_Squ_Tej
(ipTM=0.62, pTM=0.68)
Vas_Tej_Spn-E
(ipTM=0.45, pTM=0.60)
Spn-E
Tej
Vas
Spn-E
Squ
Tej
C
D
BoYb_Vret_Shu
(ipTM=0.43, pTM=0.46)
Me31B_Cup_Tral
(ipTM=0.30, pTM=0.30)
Me31B
Cup
Tral
BoYb
Vret
Shu
Supplemental Fig. S4
Trimer structures predicted by AlphaFold3
(A) SpnE_Squ_Tej.
(B) Vas_Tej_Spn-E.
(C) BoYb_Vret_Shu.
(D) Me31B_Cup_Tral.
PAE plots are also shown on the right. Orange lines indicate the protein boundaries.

### Slide 5
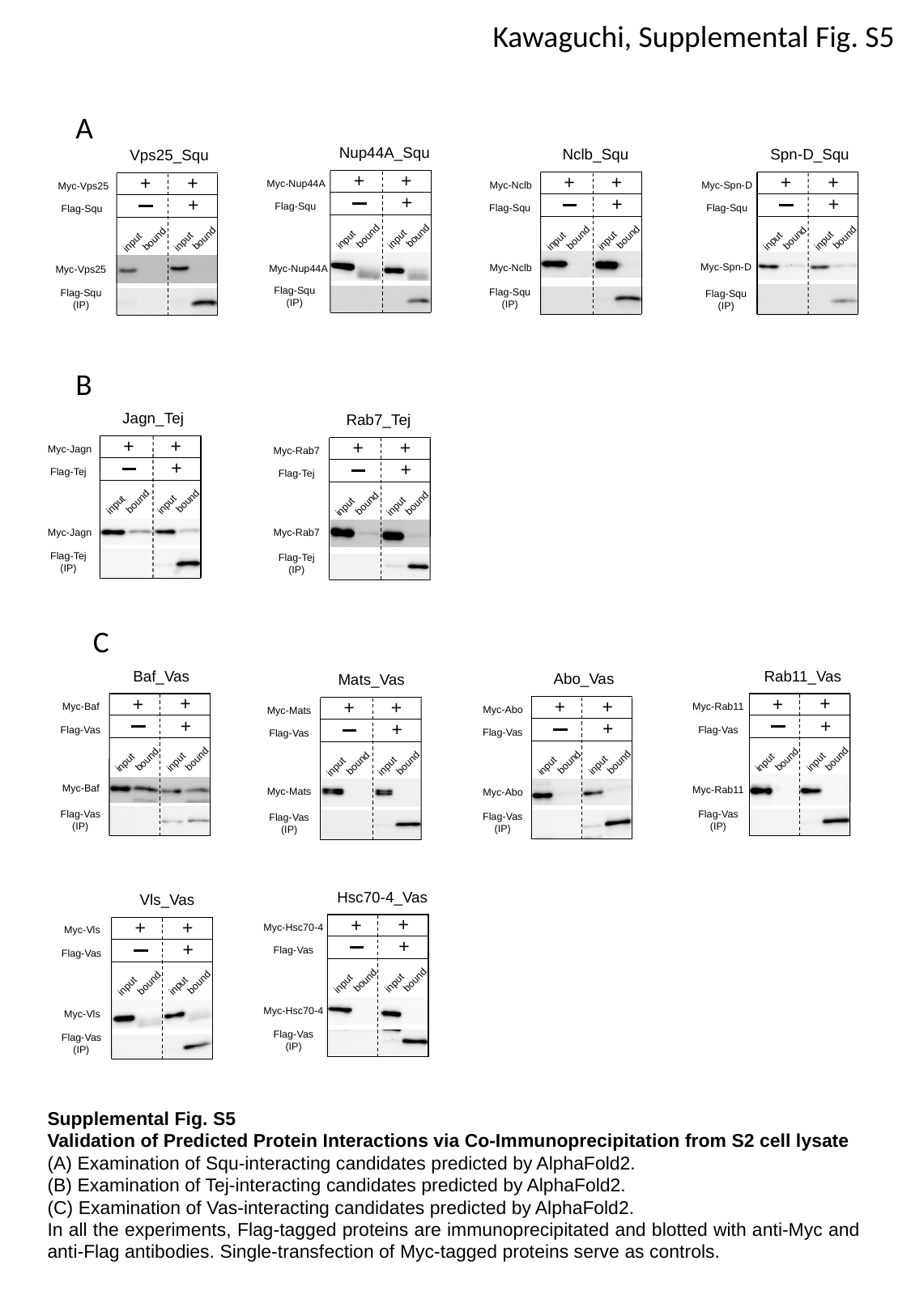

Kawaguchi, Supplemental Fig. S5
A
Nup44A_Squ
+
+
Myc-Nup44A
+
Flag-Squ
bound
bound
input
input
Myc-Nup44A
Flag-Squ
(IP)
Nclb_Squ
+
+
Myc-Nclb
+
Flag-Squ
bound
bound
input
input
Myc-Nclb
Flag-Squ
(IP)
Spn-D_Squ
+
+
Myc-Spn-D
+
Flag-Squ
bound
bound
input
input
Myc-Spn-D
Flag-Squ
(IP)
Vps25_Squ
+
+
Myc-Vps25
+
Flag-Squ
bound
bound
input
input
Myc-Vps25
Flag-Squ
(IP)
B
Jagn_Tej
+
+
Myc-Jagn
+
Flag-Tej
bound
bound
input
input
Myc-Jagn
Flag-Tej
(IP)
Rab7_Tej
+
+
Myc-Rab7
+
Flag-Tej
bound
bound
input
input
Myc-Rab7
Flag-Tej
(IP)
C
Baf_Vas
+
+
Myc-Baf
+
Flag-Vas
bound
bound
input
input
Myc-Baf
Flag-Vas
(IP)
Rab11_Vas
+
+
Myc-Rab11
+
Flag-Vas
bound
bound
input
input
Myc-Rab11
Flag-Vas
(IP)
Abo_Vas
+
+
Myc-Abo
+
Flag-Vas
bound
bound
input
input
Myc-Abo
Flag-Vas
(IP)
Mats_Vas
+
+
Myc-Mats
+
Flag-Vas
bound
bound
input
input
Myc-Mats
Flag-Vas
(IP)
Hsc70-4_Vas
+
+
Myc-Hsc70-4
+
Flag-Vas
bound
bound
input
input
Myc-Hsc70-4
Flag-Vas
(IP)
Vls_Vas
+
+
Myc-Vls
+
Flag-Vas
bound
bound
input
input
Myc-Vls
Flag-Vas
(IP)
Supplemental Fig. S5
Validation of Predicted Protein Interactions via Co-Immunoprecipitation from S2 cell lysate
(A) Examination of Squ-interacting candidates predicted by AlphaFold2.
(B) Examination of Tej-interacting candidates predicted by AlphaFold2.
(C) Examination of Vas-interacting candidates predicted by AlphaFold2.
In all the experiments, Flag-tagged proteins are immunoprecipitated and blotted with anti-Myc and anti-Flag antibodies. Single-transfection of Myc-tagged proteins serve as controls.

### Slide 6
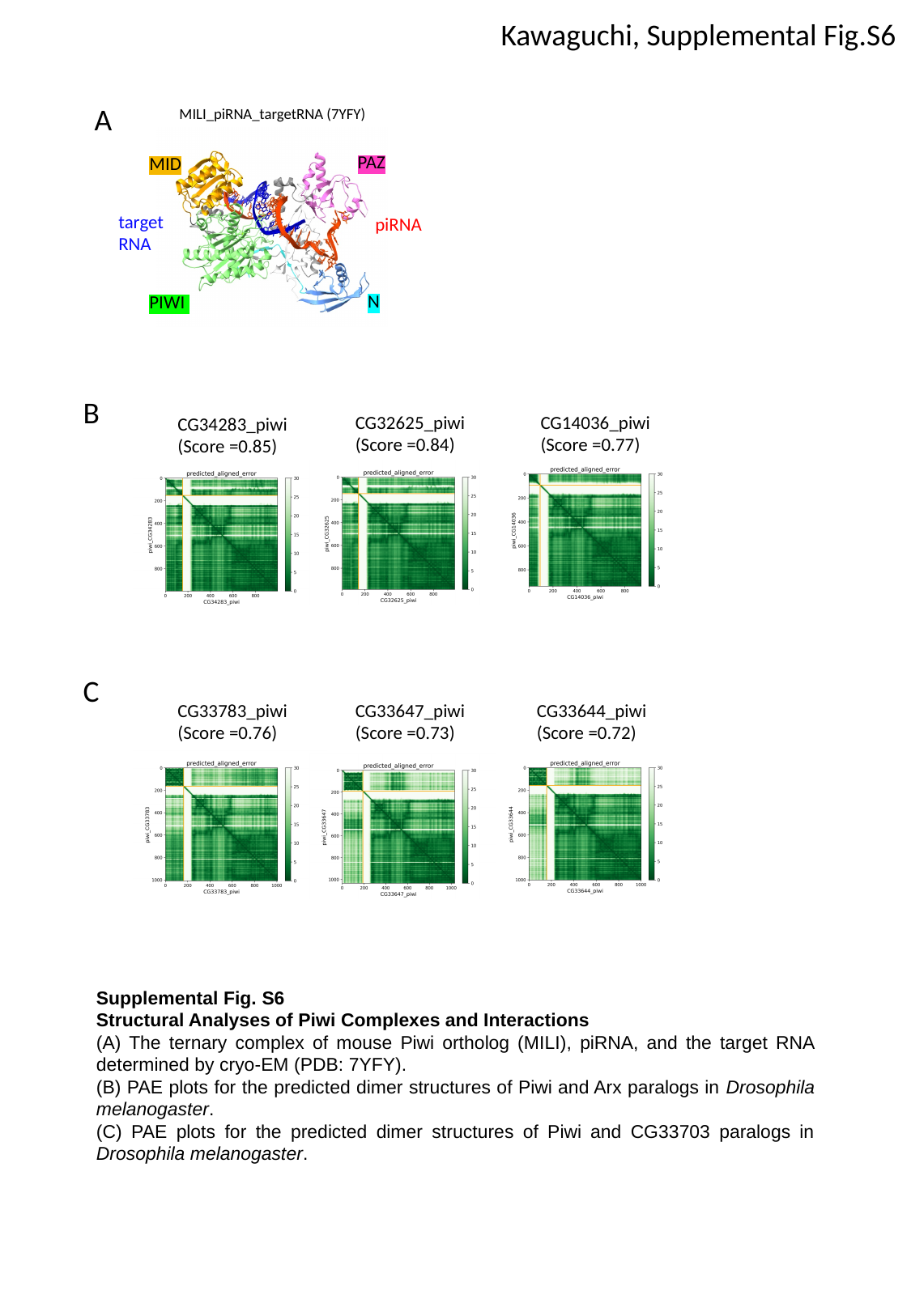

Kawaguchi, Supplemental Fig.S6
A
 MILI_piRNA_targetRNA (7YFY)
PAZ
MID
target
RNA
piRNA
N
PIWI
B
CG14036_piwi (Score =0.77)
CG32625_piwi (Score =0.84)
CG34283_piwi
(Score =0.85)
C
CG33644_piwi (Score =0.72)
CG33783_piwi (Score =0.76)
CG33647_piwi (Score =0.73)
Supplemental Fig. S6
Structural Analyses of Piwi Complexes and Interactions
(A) The ternary complex of mouse Piwi ortholog (MILI), piRNA, and the target RNA determined by cryo-EM (PDB: 7YFY).
(B) PAE plots for the predicted dimer structures of Piwi and Arx paralogs in Drosophila melanogaster.
(C) PAE plots for the predicted dimer structures of Piwi and CG33703 paralogs in Drosophila melanogaster.
