## Supplementary Table S3 for "In silico screening by AlphaFold2 program revealed the potential binding partners of nuage-localizing proteins and piRNA-related proteins"

Supplemental Table S3

The salt-bridges and H-bonds found in the predicted interface between Spn-E and Squ dimer

| **Salt bridges** |  |  |  |
| --- | --- | --- | --- |
| **#** | **Spn-E** | **Dist. [Å]** | **Squ** |
| 1 | A:ARG 788[ NH2] | 2.81 | B:GLU 103[ OE2] |
| 2 | A:LYS 834[ NZ ] | 2.79 | B:GLU 107[ OE1] |
| 3 | A:LYS 834[ NZ ] | 2.79 | B:GLU 107[ OE2] |
| 4 | A:ARG 675[ NH2] | 3.61 | B:GLU 109[ OE1] |
| 5 | A:ARG 675[ NE ] | 2.91 | B:GLU 109[ OE2] |
| 6 | A:ARG 675[ NH2] | 2.81 | B:GLU 109[ OE2] |
| 7 | A:ASP 640[ OD1] | 3.46 | B:ARG 115[ NH1] |
| 8 | A:ASP 640[ OD2] | 2.76 | B:ARG 115[ NH1] |
| 9 | A:ASP 640[ OD1] | 2.67 | B:ARG 115[ NH2] |
| 10 | A:ASP 640[ OD2] | 3.37 | B:ARG 115[ NH2] |
| 11 | A:GLU1297[ OE1] | 2.65 | B:LYS 148[ NZ ] |
| 12 | A:GLU1274[ OE1] | 3.25 | B:LYS 163[ NZ ] |
| 13 | A:GLU1274[ OE2] | 2.70 | B:LYS 163[ NZ ] |
| **Hydrogen bonds** |  |  |  |
| **#** | **Spn-E** | **Dist. [Å]** | **Squ** |
| 1 | A:ARG 675[HH12] | 2.44 | B:ASP  99[ O  ] |
| 2 | A:ARG 675[HH11] | 2.40 | B:TRP 100[ O  ] |
| 3 | A:ARG 675[HH12] | 1.93 | B:ASP 102[ O  ] |
| 4 | A:ARG 675[HH22] | 2.01 | B:ASP 102[ O  ] |
| 5 | A:ARG 788[HH11] | 1.94 | B:GLU 103[ O  ] |
| 6 | A:ARG 788[HH21] | 1.93 | B:GLU 103[ OE2] |
| 7 | A:LYS 834[ HZ3] | 1.80 | B:GLU 107[ OE2] |
| 8 | A:ARG 675[ HE ] | 1.98 | B:GLU 109[ OE2] |
| 9 | A:ARG 675[HH21] | 1.88 | B:GLU 109[ OE2] |
| 10 | A:ARG 651[HH11] | 1.77 | B:MET 120[ O  ] |
| 11 | A:ARG1028[HH12] | 1.87 | B:ALA 123[ O  ] |
| 12 | A:ASN1281[HD22] | 2.49 | B:PHE 159[ O  ] |
| 13 | A:LEU1364[ H  ] | 2.00 | B:ARG 162[ O  ] |
| 14 | A:GLU1366[ H  ] | 1.85 | B:ILE 164[ O  ] |
| 15 | A:ASP 665[ OD2] | 2.06 | B:GLN  74[HE22] |
| 16 | A:ASP 640[ OD2] | 1.77 | B:ARG 115[HH11] |
| 17 | A:ASP 640[ OD1] | 1.66 | B:ARG 115[HH22] |
| 18 | A:SER1309[ OG ] | 1.96 | B:LYS 141[ HZ1] |
| 19 | A:GLU1306[ OE1] | 2.21 | B:ARG 144[ H  ] |
| 20 | A:ALA1301[ O  ] | 1.91 | B:ASN 145[HD22] |
| 21 | A:GLU1297[ OE1] | 1.65 | B:LYS 148[ HZ3] |
| 22 | A:TYR1285[ OH ] | 1.97 | B:ILE 153[ H  ] |
| 23 | A:PRO1347[ O  ] | 2.03 | B:ARG 162[HH12] |
| 24 | A:GLU1274[ OE2] | 1.75 | B:LYS 163[ HZ2] |
| 25 | A:LEU1364[ O  ] | 1.89 | B:ILE 164[ H  ] |
| 26 | A:LYS1236[ O  ] | 2.17 | B:ARG 166[HH11] |
| 27 | A:ASN1235[ O  ] | 2.07 | B:ARG 166[HH22] |
